# Genetically Encoded RNA-based Bioluminescence Resonance Energy Transfer (BRET) Sensors

**DOI:** 10.1101/2022.09.28.509942

**Authors:** Lan Mi, Qikun Yu, Aruni P.K.K. Karunanayake Mudiyanselage, Rigumula Wu, Zhining Sun, Ru Zheng, Kewei Ren, Mingxu You

## Abstract

RNA-based nanostructures and molecular devices have become popular for developing biosensors and genetic regulators. These programmable RNA nanodevices can be genetically encoded and modularly engineered to detect various cellular targets and then induce output signals, most often a fluorescence readout. Although powerful, the high reliance of fluorescence on the external excitation light raises concerns on its high background, photobleaching, and phototoxicity. Bioluminescence signals can be an ideal complementary readout for these genetically encoded RNA nanodevices. However, RNA-based real-time bioluminescent reporters have been rarely developed. In this study, we reported the first type of genetically encoded RNA-based bioluminescence resonance energy transfer (BRET) sensors that can be used for real-time target detection in living cells. By coupling a luciferase bioluminescence donor with a fluorogenic RNA-based acceptor, our BRET system can be modularly designed to image and detect various cellular analytes. We expect this novel RNA-based bioluminescent system can be potentially used broadly in bioanalysis and nanomedicine for engineering biosensors, characterizing cellular RNA–protein interactions, as well as high-throughput screening or *in vivo* imaging.

## Introduction

Genetically encodable fluorescent and bioluminescent sensors are most widely used tools for the live-cell imaging and detection of various biomolecules and signaling events.^1-3^ While fluorescent molecules rely on the absorption of excitation light to reemit photons, bioluminescence can produce light directly through chemical reactions. Compared to bioluminescence, although fluorescence signals are often brighter, the requirement of excitation light may cause higher background, photobleaching, and phototoxicity.^4,5^ Bioluminescence and fluorescence have thus been frequently used as complementary tools for studying cellular processes.

A wide variety of protein-based fluorescent and bioluminescent sensors have been developed for both cellular and *in vivo* studies.^5-9^ In contrast, only recently, RNA-based genetically encodable fluorescent sensors began to be engineered for intracellular measurement.^10^ The major advantages of these RNA-based sensors include their high sensitivity, modularity, and ease of engineering and evolution. As a result, different RNAs, proteins, and small-molecule analytes can be detected and imaged by these RNA-based fluorescent sensors.^11-14^ On the other hand, RNA-based bioluminescent sensors remain largely underdeveloped. RNA-based genetic regulators can induce bioluminescence signals by controlling the cellular expression of luciferase reporters.^2,11,15^ However, these RNA nanodevices can rarely be used as biosensors for the real-time detection or monitoring of cellular target analytes.

In this project, our goal is to design a novel type of genetically encoded RNA biosensors that can produce real-time bioluminescence signals inside living cells, without the use of excitation light. Our strategy is based on the modulation of bioluminescence resonance energy transfer (BRET) between a donor luciferase enzyme and an acceptor fluorescent RNA species (Fig. 1a). An important advantage of BRET-based biosensors is that they can provide ratiometric signals to allow real quantitative measurement of target analytes.^7,16^

**Fig. 1.**
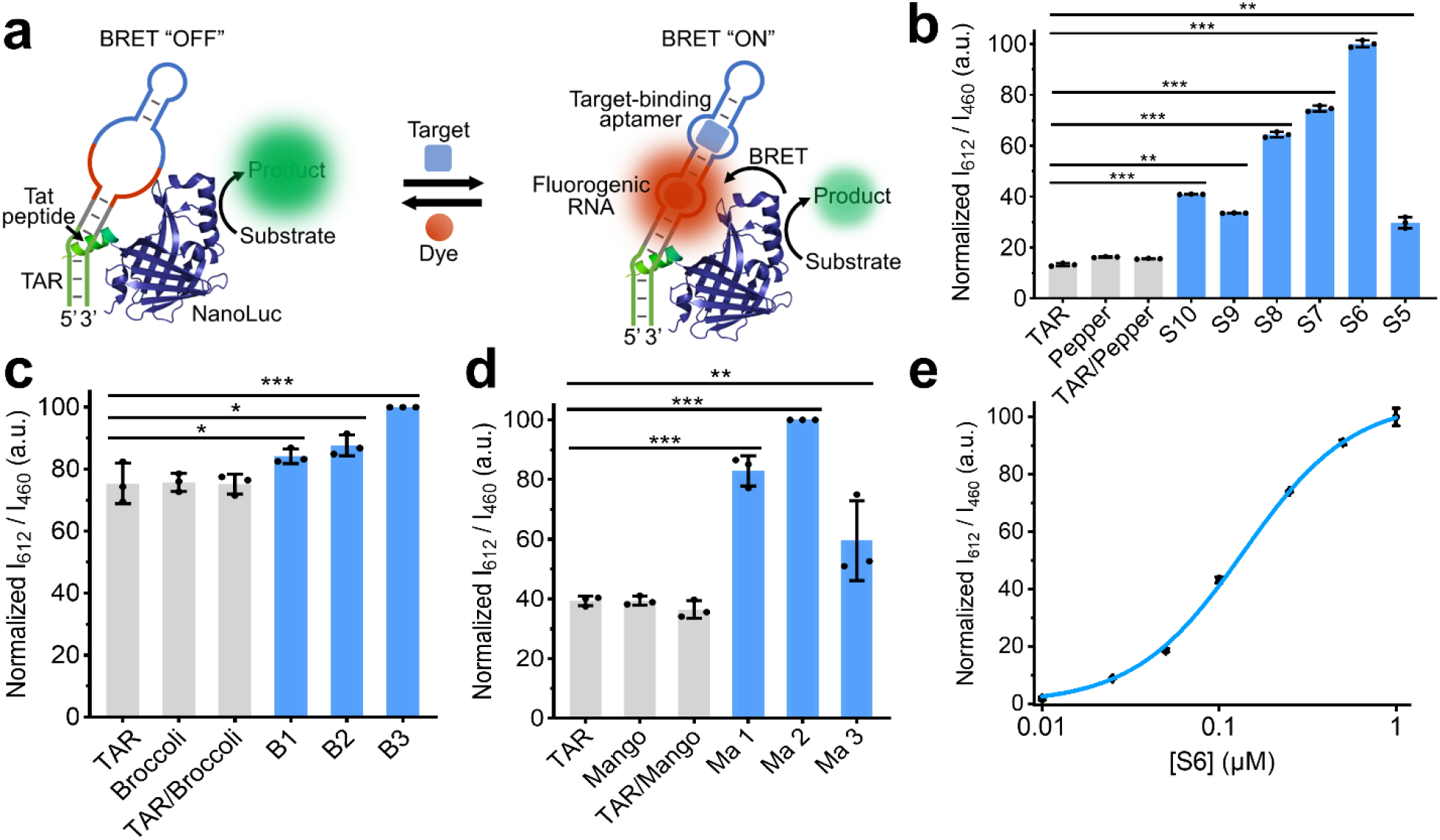
(**a**) Schematic illustration of a modular BRET sensor system based on TAR–Tat interaction-induced energy transfer between NanoLuc/substrate complex and fluorogenic RNA/dye pairs. The target binding to the aptamer region induces the folding of fluorogenic RNA to bind the dye molecule. (**b**) The I_612_/I_460_ (acceptor/donor) ratiometric BRET signal was measured at 25°C in a solution containing 100 nM t-NLuc, 1 μM RNA, 5 μM HBC620, and 5 mM Mg^2+^ at 30 s after adding 1% (v/v) furimazine substrate. S5–S10 represent the t-NLuc/TAR-Pepper construct containing 5–10-base-pair-long linkers, respectively. The mixture of t-NLuc with TAR, Pepper, and their simple mixture (TAR/Pepper) acted as the negative controls. (**c, d**) The BRET efficiency of Broccoli- and Mango II-based t-NLuc/RNA construct with different linker lengths (named as B1, B2, B3 and Ma1, Ma2, Ma3). The I_507_/I_460_ and I_535_/I_460_ ratiometric signals were measured in a solution containing 100 nM t-NLuc and 1 μM RNA at 25°C, in the presence of 20 μM DFHBI-1T or 1 μM TO1-biotin. The mixture of t-NLuc with TAR (without Broccoli or Mango II), Broccoli, Mango II (without TAR conjugation), or simple mixtures of Broccoli/Mango II and TAR (i.e., TAR/Broccoli and TAR/Mango) were used as negative controls. (**e**) The normalized I_612_/I_460_ BRET signal of the t-NLuc/S6 construct after mixing 100 nM t-NLuc with different concentrations of the S6 RNA. Shown are the mean and standard deviation (SD) values from three independent replicates. *p< 0.05, **p< 0.01, ***p< 0.001 in two-tailed student’s t-test.

Several protein-based BRET sensors have been previously developed to detect different cellular target molecules or protein–protein interactions.^2,17,18^ In these sensors, the ratiometric BRET signals are generated between an adjacent pair of luciferase and fluorescent protein. The brightness and stability of the luciferase and substrate is critical to the sensitivity of the BRET sensors.^18,19^ For example, the NanoLuc luciferase (**NLuc**) has been popularly used as a BRET donor due to its small size, high catalytic efficiency and stability.^20,21^

To develop an RNA-based BRET system, herein, we chose to continue using luciferase (e.g., NLuc) as the donor moieties, but will replace the fluorescent protein acceptors with fluorogenic RNA (**FR**) aptamers. FR aptamers are single-stranded RNAs that can bind and activate the fluorescence of corresponding small-molecule dyes.^22,23^ We and others have previously demonstrated that these FRs can be engineered into modular fluorescent sensors for the sensitive imaging of various cellular targets.^11-14,24-29^ We speculate that once our RNA-based BRET platform was established, these existing fluorescent RNA sensors can be readily converted to generate target-specific bioluminescence signals.

Herein, we will report the design and validation of the first RNA-based genetically encodable BRET system based on natural RNA–protein interactions. We have further incorporated modular FR sensors to engineer bioluminescent probes for the detection of different small-molecule analytes inside living cells. With the development of this new type of RNA-based BRET sensors, a broad range of cellular target analytes, as well as RNA–protein interactions, can be potentially imaged and quantified *in vitro*, or even *in vivo*.

## Materials and methods

### Chemicals and reagents

All the chemicals were purchased from Sigma or Fisher Scientific unless otherwise noted. Furimazine (Nano-Glo^®^ Luciferase Assay) was purchased from Promega (Madison, WI). All the RNA structures were simulated and designed using an Mfold online software, and all the RNA sequences used in this project are listed in Table S1. DNA oligonucleotides were synthesized and purified by W.M. Keck Oligonucleotide Synthesis Facility (Yale University School of Medicine) or Integrated DNA Technologies (Coralville, IA). The stocks were dissolved at 100 μM concentration in 10 mM Tris-HCl, 0.1 mM EDTA at pH=7.5 and stored at –20°C. Double-stranded DNA template/non-template strands for *in vitro* transcription were prepared via PCR amplification using an Eppendorf Mastercycler. The PCR products were purified and cleaned using a Monarch^®^ PCR & DNA cleanup kit (New England BioLabs, Ipswich, MA). RNAs for the *in vitro* test were transcribed using a HiScribe™ T7 high yield RNA synthesis kit (New England BioLabs, Ipswich, MA) and then column purified. These RNA strands were prepared in aliquots and stored at –20°C for immediate usage or at –80°C for long-term storage. All the DNA/RNA concentrations were measured using a NanoDrop One UV-vis spectrophotometer.

### Bioluminescence and fluorescence measurement

All the *in vitro* bioluminescence and fluorescence measurements were conducted using a PTI fluorometer (Horiba, New Jersey, NJ) or a SpectraMax^®^ M2 multimode microplate reader. Fluorescence assays were conducted in a home-made buffer consisting of 40 mM HEPES, 100 mM KCl, 0.1% DMSO and 1–5 mM MgCl_2_, at pH=7.5. Here, 1 μM RNA and 5 μM HBC620 (or 20 μM DFHBI-1T for Broccoli, or 1 μM TO1-biotin for Mango II) was used for these measurements at 25°C. Bioluminescence assays were performed either in the same home-made buffer or a NanoBuffer™ (Promega Nano-Glo Luciferase Assay), with the addition of 100 nM t-NLuc, 1 μM RNA and their corresponding dye. All these samples were first mixed and incubated at 25°C for 30 min before taking the measurement. Right after the addition of 1% (v/v) furimazine, the BRET signals of each sample were collected at 460 nm (donor) and 612 nm (Pepper acceptor), 507 nm (Broccoli acceptor), or 535 nm (Mango II acceptor) without excitation. All the analyzed data were plotted using an Origin software.

### Vector construction

The DNA sequence of the t-NLuc was first PCR amplified from a miniCMV-NLuc-tDeg plasmid^15^ (gift from the Jaffrey Lab). The amplified t-NLuc sequence was then cloned into a pETite vector containing His tag for protein purification and *in vitro* characterization. For the intracellular bioluminescent detection, sequences containing both a T7 promoter and terminator were cloned into a pETDuet vector. The t-NLuc-His tag was double-digested with NdeI and XhoI restriction enzymes (New England BioLabs), and the RNA sensor region were digested with SgrAI and SacII restriction enzymes. After gel purification, the digested vector was ligated with a similarly digested DNA insert using a T4 DNA ligase (New England BioLabs). The preparation of t-NLuc under the lac promoter in the reconstructed pETDuet plasmid was achieved following a similar procedure but was double-digested with SgrAI and SacII restriction enzymes. The ligated product was transformed into BL21 Star™ (DE3) cells (New England Biolabs) and screened based on their ampicillin resistance. All these plasmids were isolated and confirmed by Sanger sequencing at Eurofins Genomics.

### BRET measurement in bacterial cells

BL21 Star™ (DE3) cells that express t-NLuc and RNA sensors were first grown in LB media at 37°C until the optical density at 600 nm reaching 0.4–0.5. Then 1 mM isopropyl β-d-1-thiogalactopyranoside (IPTG) was added for a 2-hour induction. Afterwards, 1 mL cell was centrifuged down for 2 min at 5000x g speed and then resuspended in 200 μL Dulbecco′s phosphate buffered saline (DPBS) or cell lysis buffer containing 5 μM HBC620. Target analyte was added into 100 μL cell solutions and incubated for 30 min before adding furimazine. The bioluminescence and fluorescence signals were collected within a 96-well plate using either a SpectraMax^®^ M2 multimode microplate reader or an IVIS^®^ SpectrumCT system. On the plate reader, the BRET signals were collected at both 460 nm and 612 nm wavelength without excitation for 3 times with calibration at 25°C. All the data were plotted with an Origin software. In the IVIS^®^ SpectrumCT system, the BRET signals were collected at both 500 nm and 620 nm wavelength without excitation. Images were taken with a 0.2 second exposure time, 1.5 cm subject height, and 13.3 cm^2^ field of view. The processing of all images was accomplished using a Fiji ImageJ software.

### BRET measurement in mammalian cells

HEK-293T cells were cultured in a DMEM medium supplemented with 10% fetal bovine serum, 100 unit/mL penicillin, and 0.1 mg/mL streptomycin at 37°C in a 5% CO_2_ atmosphere. For the microscope imaging, HEK-293T cells were seeded at a density of 2×10^5^ cells per well into poly-D-lysine-coated 8 chambered cover glass plate (Cellvis, C8-1.5H-N). For the plate reader experiment, these HEK-293T cells were seeded at a density of 5×10^4^ cells per well into 96-well glass bottom plate (Cellvis, P96-1.5H-N). After overnight culturing, cells were transfected using the FuGENE^®^ HD transfection reagent (Promega) according to the manufacturer’s instructions. To be more specific, 1 μg mini-CMV-t-NLuc was co-transfected with 1 μg PAV-U6+27-Tornado-TAR or 1 μg PAV-U6+27-Tornado-S6. After 24 h of transfection, cells were sub-cultured using a DPBS buffer containing 5 μM HBC620 for 30 min. Fluorescence images were collected using a Nikon TiE inverted microscope with an Andor’s Zyla sCOMS camera at 25°C. Cells were excited with a 561 nm laser through a 40x oil objective. For the plate reader experiment, furimazine was added directly in each well before reading, the donor luminescence signal at 460 nm and the acceptor signal at 645 nm were collected using a BioTek Synergy 2 plate reader.

## Results and discussion

### Design and optimization of an RNA-based BRET system

Inspired by previously reported BRET sensors that are regulated by protein–protein interactions,^7,30-33^ we first wondered if natural RNA–protein complexes can also be used to control BRET signals. To test this idea, we chose to study a well characterized bovine immunodeficiency virus interaction between a trans-activation response (**TAR**) RNA and an arginine-rich **Tat** peptide.^34-36^ This TAR–Tat complex exhibits strong binding affinity (K_D_, ∼10 nM) and is naturally used to enhance transcriptional activation and elongation.^37-39^ Guided by the structure of this RNA–peptide complex,^40,41^ we fused an NLuc luciferase to the N-terminus of the Tat peptide (named as “**t-NLuc**”) and used it as the BRET donor (Fig. 1a). Three types of FR aptamers, i.e., Broccoli,^42^ Mango II,^43,44^ and Pepper,^45,46^ were respectively conjugated to the loop region of TAR and acted as the potential BRET acceptor (Table S1). This loop region of TAR has been shown to just play a structural role, and thus can be swapped into different sequences without interfering the formation of the TAR–Tat complex.^41,47^ The three types of FRs were chosen based on their high brightness upon activating the corresponding dyes, i.e., DFHBI-1T, TO1-biotin, and HBC620. These FR/dye pairs can also cover a large spectral range, i.e., Broccoli/DFHBI-1T (λ_ex_/λ_em_, 472/507 nm), Mango II/TO1-biotin (λ_ex_/λ_em_, 510/535 nm), and Pepper/HBC620 (λ_ex_/λ_em_, 572/612 nm).

Because the BRET efficiency strongly depends on the distance and orientation between the donor and acceptor modules, we next wanted to optimize these factors between NLuc and FRs. Instead of changing the peptide linker that connects Tat with NLuc, it is much easier to synthesize and adjust the RNA linker between TAR and FR aptamers. By simply altering the length of RNA linker, considering the A-form helical conformation of RNA duplex, we can fine-tune both the distance and orientation between NLuc and FRs. For example, in the case of the Pepper acceptor, 5–10-base-pair-long linkers (named as **S5–S10**, respectively) were synthesized and tested *in vitro* (Table S1). Our results indicated that the fusion of Pepper and TAR via these linkers won’t influence the Pepper/HBC620 fluorescence signal (Fig. S1a). While in the presence of t-NLuc, upon adding furimazine, the NanoLuc substrate, the BRET signals can indeed be observed but highly dependent on the length of RNA linkers (Fig. S1b).

To compare the efficiency among these different BRET pairs, the bioluminescence ratios between the acceptor (e.g., I_612_ for Pepper) and donor (I_460_) channels were determined.^20,45^ Measurement of these ratiometric I_612_/I_460_ signals can minimize the influence of incubation time and substrate concentration on the bioluminescence readout (Fig. S2). Our results showed that in the presence of S5–S10 RNA, a 1.2-to 7.2-fold increase in the I_612_/I_460_ BRET signals was observed (Fig. 1b). The maximum I_612_/I_460_ ratiometric signal was detected with the **t-NLuc/S6** complex. As a control, without linking TAR with Pepper, a minimal I_612_/I_460_ signal was shown, indicating that the TAR– Tat interaction is required for the generation of strong BRET signals.

We have also measured the BRET efficiency of other t-NLuc/FR system by using Broccoli/DFHBI-1T or Mango II/TO1-biotin as the acceptor. After optimizing the linker lengths between TAR and FR (Table S1), the ratiometric BRET signals were determined at I507/I460 and I535/I460, respectively. However, only up to 33% and 156% increase in the BRET signal was observed in the presence of Broccoli or Mango II acceptor, respectively (Fig. 1c and 1d). Although these results indicated that different types of FRs can be potentially used to generate bioluminescent signals, considering the much larger signal enhancement and red-shifted emission spectra from the Pepper/HBC620 system, we decided to use the t-NLuc/S6 platform for the following studies.

To further optimize the BRET signals of this t-NLuc/S6 system, we studied the effect of Mg^2+^ concentration (Fig. S3) and other buffer conditions (Fig. S4). Magnesium ions can potentially influence the binding affinity of the TAR–Tat complex^47^ as well as the folding of Pepper RNA.^45^ Indeed, without adding Mg^2+^, a minimal I_612_/I_460_ signal was observed (Fig. S3). While at physiological Mg^2+^ level (∼2 mM),^48,49^ a ∼11.0-fold increase in the I_612_/I_460_ BRET signal was shown, indicating that t-NLuc/S6 may be potentially used under normal biological conditions. As we further increased the Mg^2+^ concentration from 2 mM to 20 mM, a gradual decrease in the I612/I460 signal was detected, which may be due to the formation of other unwanted RNA tertiary structures.

In addition, we wondered if these BRET signals are indeed correlated with the concentrations of S6 RNA. To test this, we mixed 100 nM t-NLuc with different amounts of S6 and measured their I612/I460 ratiometric signals (Fig. 1e). Our results indicated that a large concentration range (10–500 nM) of S6 can be distinguished based on their BRET signals. A half-maximal ratiometric intensity was reached after adding ∼100 nM S6, i.e., stoichiometrically equivalent to that of t-NLuc.

### Development of RNA-based BRET sensors

After characterizing and optimizing the function of the t-NLuc/S6 system, we next asked if this platform can be further used to develop sensors for detecting target analytes. Inspired by the modular design of FR-based allosteric fluorescent sensors,^22^ we wondered if RNA-based BRET sensors can also be engineered similarly by fusing target-binding aptamers into the S6 RNA (Fig. 1a). In this case, upon the addition of target analytes, target-binding aptamers undergo structural changes to regulate the folding of Pepper RNA, i.e., the BRET acceptor. Consequently, the I_612_/I_460_ ratiometric BRET signal will be directly correlated with the amount of target analytes.

We realized that the P2 stem region of Pepper is sequence-independent and thus can be potentially used to insert these target-binding aptamers.^46^ To test this design principle, we first developed BRET sensors for detecting tetracycline, a widely used antibiotics for fighting bacterial infections.^50^ Five different double-stranded transducers were designed to link the tetracycline aptamer into the S6 (Fig. 2a and Table S1). These transducers were chosen because they have been previously applied to engineer modular RNA-based tetracycline sensors.^51^ Each of these sensor structures was further characterized *in silico* using a Mfold online software.^52^ By measuring the I_612_/I_460_ BRET signal of each RNA sensor in the presence or absence of 100 μM tetracycline using a solution containing 1 μM RNA and 100 nM t-NLuc, an optimal sensor (**T1**) was identified that exhibited a ∼3.4-fold increase in the ratiometric BRET signal (Fig. 2a and S5a).

**Fig. 2.**
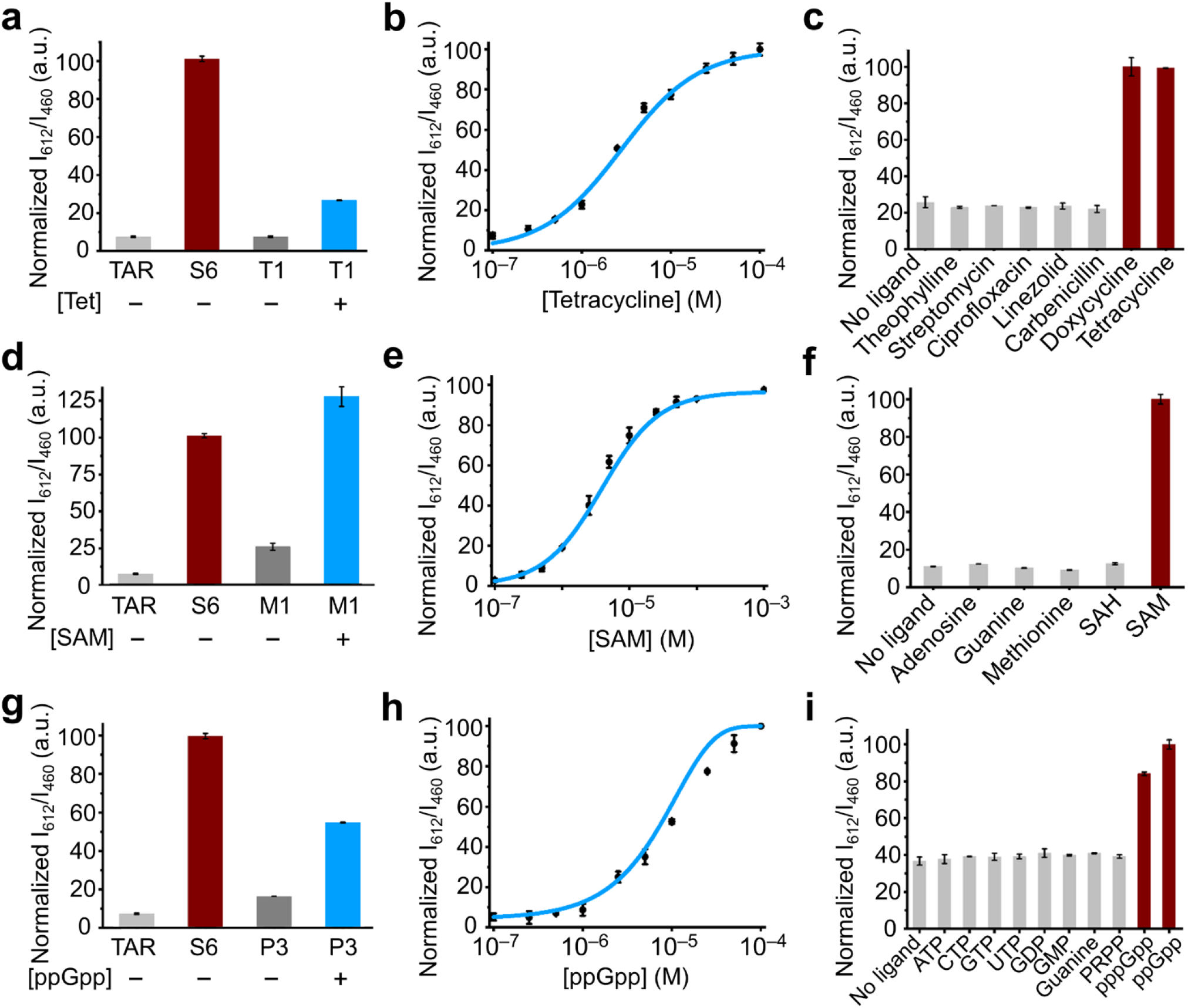
In vitro characterization of different RNA-based BRET sensors. The I_612_/I_460_ ratiometric signals of the optimal (**a**) t-NLuc/T1 tetracycline sensor, (**d**) t-NLuc/M1 SAM sensor and (**g**) t-NLuc/P3 ppGpp sensor were measured in the presence or absence of 100 μM tetracycline (Tet), SAM or ppGpp. The mixture of t-NLuc with TAR or S6 acted as a negative control and positive control, respectively. (**b, e, h**) Dose response curve of the t-NLuc/T1 tetracycline sensor, t-NLuc/M1 SAM sensor and t-NLuc/P3 ppGpp sensor after adding different amounts of tetracycline, SAM or ppGpp. (**c, f, i**) Selectivity of the t-NLuc/T1 tetracycline sensor, t-NLuc/M1 SAM sensor and t-NLuc/P3 ppGpp sensor was measured in a solution containing 100 μM counter ligands. Control was measured without adding ligands. All these measurements were performed in a solution containing 100 nM t-NLuc, 1 μM RNA, and 5 μM HBC620 at 25°C after adding 1% (v/v) furimazine substrate. Shown are the mean and the standard error of the mean values from three independent replicates in each case.

A dose-response curve for tetracycline was further measured using this t-NLuc/T1 sensor (Fig. 2b). The EC_50_ value of this tetracycline-targeting BRET system was determined to be ∼2.8 μM. The t-NLuc/T1 sensor can be used for detecting a large concentration range (0.1–100 μM) of tetracycline.

As a control, the bioluminescent signals from either t-NLuc/TAR or t-NLuc/S6 were not affected by incubating with 100 μM of tetracycline (Fig. S5b). To further test the target selectivity of this BRET sensor, we incubated 100 μM theophylline, streptomycin, ciprofloxacin, doxycycline, linezolid, or carbenicillin with this t-NLuc/T1 sensor system (Fig. 2c). As expected, minimal I_612_/I_460_ signal was shown, except for doxycycline, which belongs to the same tetracycline family and shares a quite similar four-ring structure as tetracycline.^53^

We wondered if this new RNA-based BRET sensor design can also be modularly used to develop sensors for different target analytes, such as S-adenosyl methionine (SAM), metabolite for methylation, transsulfuration, polyamine synthesis,^54,55^ and guanosine tetraphosphate (ppGpp), signaling molecule involved in bacterial response to stringent conditions.^56-58^ Naturally existing RNA riboswitches that can selectively recognize SAM^59^ and ppGpp^60,61^ have been identified and used previously for engineering FR-based fluorescent sensors.^13,62^

To convert these riboswitches into BRET sensors, we again focused on the design of transducer sequences between the aptamer domain of riboswitch and the Pepper P2 stem region (Fig. 1a and Table S1). In the presence of t-NLuc, after mixing these candidate RNA sensors with 100 μM SAM or ppGpp, a 10.0-fold and 2.2-fold enhancement in the I_612_/I_460_ ratiometric signal was observed for the optimal SAM (t-NLuc/**M1**) and ppGpp (t-NLuc/**P3**) sensors, respectively (Fig. 2d, 2g, and S6). While as expected, the I_612_/I_460_ signals of t-NLuc/TAR and t-NLuc/S6 controls were not affected after adding 100 μM SAM or ppGpp (Fig. S5). These optimized BRET sensors can be used to detect 1–100 μM SAM and ppGpp, with an EC_50_ value of 5.2 μM and 7.1 μM, respectively (Fig. 2e and 2h).

We next tested if these BRET sensors can retain the target selectivity of natural riboswitches. For this purpose, we incubated adenosine, guanine, methionine, and S-adenosylhomocysteine (SAH) with the t-NLuc/M1 sensor, and ATP, CTP, GTP, UTP, GDP, GMP and guanine with the t-NLuc/P3 sensor. None of these compounds affected the I_612_/I_460_ signal of the BRET sensors (Fig. 2f and 2i). Indeed, the target selectivity of the riboswitches is maintained in these t-NLuc/RNA-based BRET sensors. All together, these above results indicated that by simply changing the aptamer domain, RNA-based BRET sensors can be modularly developed to allow the detection of various target analytes.

### Live-cell detection with genetically encoded RNA-based BRET sensors

We next asked if these t-NLuc/S6-based BRET sensors can be genetically encoded for intracellular measurement. We first cloned S6 and t-NLuc into a pETDuet vector to allow the co-expression of S6 RNA and t-NLuc protein under two separate T7 promoters. As a control, another pETDuet vector was engineered to express only the t-NLuc. After transforming these two plasmids respectively into BL21 Star™ (DE3) *E. coli* cells and incubating with HBC620 and furimazine, a strong donor bioluminescent signal at ∼460 nm was clearly detected (Fig. S6). However, minimal I612 acceptor signal was shown in these t-NLuc/S6-expressing cells.

To solve this problem, we tried to change the relative cellular expression levels of S6 and t-NLuc. This is because as shown in Fig. 1c, an enhanced BRET efficiency can be observed with an increasing ratio of S6:t-NLuc. We thus decided to suppress the cellular expression of t-NLuc by placing it under a weaker lac promotor (in substitution of T7), while S6 was continued to be expressed under a strong T7 promotor. This new expression system allows the generation of an increased S6:t-NLuc concentration ratio inside cells, and indeed in this case, much clearer cellular I_612_/I_460_ BRET signals were observed (Fig. S6).

Our next goal was to study the performance of RNA-based BRET sensors inside these living *E. coli* cells. Again, we engineered the vectors to co-express lac promotor-regulated t-NLuc and T7 promoter-controlled RNA sensors. Vectors expressing t-NLuc/T1 tetracycline sensor, t-NLuc/M1 SAM sensor, and t-NLuc/P3 (p)ppGpp sensor were separately prepared, and then respectively transformed into BL21 Star™ (DE3) *E. coli* cells. We first tried to detect the cellular accumulation of tetracycline. In this experiment, after adding 100 μM tetracycline, a ∼3.4-fold enhancement in the I612/I460 BRET signal was detected in the t-NLuc/T1-expressing cells (Fig. 3a). The addition of 0.1– 100 μM tetracycline can be distinguished based on the cellular BRET signals. This detection range was similar to that from *in vitro* characterizations (Fig. 2b).

**Fig. 3.**
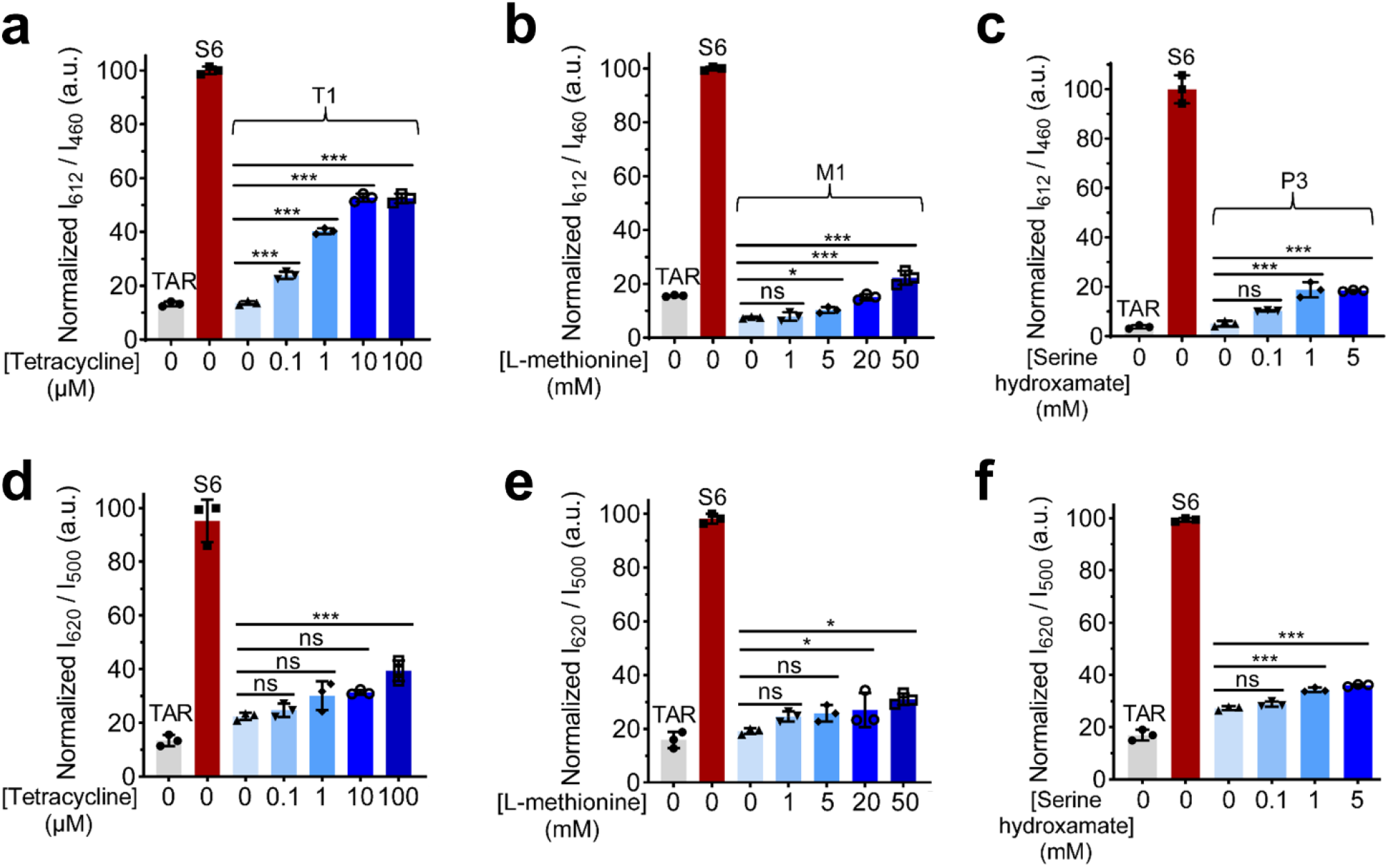
Target detection in living BL21 Star™ (DE3) *E. coli* cells with different RNA-based BRET sensors. The ratiometric signals of t-NLuc/T1 sensor-expressing cells in the presence of different amounts of tetracycline using (**a**) plate reader (I_612_/I_460_) and (**d**) IVIS^®^ SpectrumCT (I_620_/I_500_). The ratiometric signals of t-NLuc/M1 sensor-expressing cells in the presence of different amounts of L-methionine, a precursor for the cellular synthesis of SAM, using (**b**) plate reader (I_612_/I_460_) and (**e**) IVIS^®^ SpectrumCT (I_620_/I_500_). The ratiometric signals of t-NLuc/P3 sensor-expressing cells in the presence of different amounts of serine hydroxamate (SHX) for inducing bacterial starvation and ppGpp generation, using (**c**) plate reader (I_612_/I_460_) and (**f**) IVIS^®^ SpectrumCT (I_620_/I_500_). Cells expressing t-NLuc/TAR or t-NLuc/S6 acted as a negative control and positive control, respectively. The measurement was performed at 25°C in the presence of 5 μM HBC620 right after adding 1% (v/v) furimazine substrate. Shown are the mean and SD values from three independent replicates. *p< 0.05, ***p< 0.001 in two-tailed student’s t-test; ns, not significant.

We next asked if the SAM- and ppGpp-targeting BRET sensors can indeed be used to detect these cellular endogenous analytes. We first treated the t-NLuc/M1-expressing *E. coli* cells with 1– 50 mM of L-methionine, a precursor that facilitates the cellular synthesis of SAM.^63^ Upon the addition of L-methionine, an up to 2.0-fold increase in the cellular BRET signal was observed (Fig. 3b). We also applied the t-NLuc/P3-expressing BL21 Star™ (DE3) cells to measure variations in the cellular ppGpp levels under different nutritional conditions. These sensor-expressing *E. coli* cells were either cultured in a nutrient-rich medium containing casamino acids and glucose^64^ or a nutrient-poor M9 minimal medium supplemented with serine hydroxamate to induce bacterial starvation.^65^ Indeed, in the nutrient-poor condition, an around 2.0-fold increase in the BRET signal was observed, indicating the cellular generation of ppGpp as a stringent response (Fig. 3c). All these data validated that these RNA-based BRET sensors can be used for detecting target analytes in living cells.

Moreover, we further tested if these t-NLuc/RNA-based BRET signals are bright enough to be detectable using an IVIS^®^ SpectrumCT system, a widely used preclinical *in vivo* study instrument. Similar to the above bioluminescence results (based on plate reader), target-induced BRET signals can be clearly detected using the IVIS^®^ SpectrumCT (Fig. 3d–f). A 26%–75% enhancement in the cellular ratiometric signals was shown after adding 100 μM tetracycline, 50 mM L-methionine, or 5 mM serine hydroxamate. These results indicated that these RNA-based BRET sensors have sufficient brightness that may be potentially used for *in vitro* or even *in vivo* studies of cellular target analytes.

It is also worth mentioning that these RNA-based BRET sensors can also function in cell lysates. For example, in the cell lysates of t-NLuc/T1-, t-NLuc/M1-, or t-NLuc/P3-expressing BL21 Star™ (DE3) cells, upon the addition of 100 μM tetracycline, 1 mM SAM, or 100 μM ppGpp, an approximately 119%, 21%, and 90% enhancement in the I_612_/I_460_ ratiometric signals was observed (Fig. 4a–4c). Interestingly, the performance of these RNA-based BRET sensors in living bacterial cells are even superior to that in the cell lysates, which is probably due to the influence of cell lysates on the RNA folding and/or the strength of TAR–Tat interactions.

**Fig. 4.**
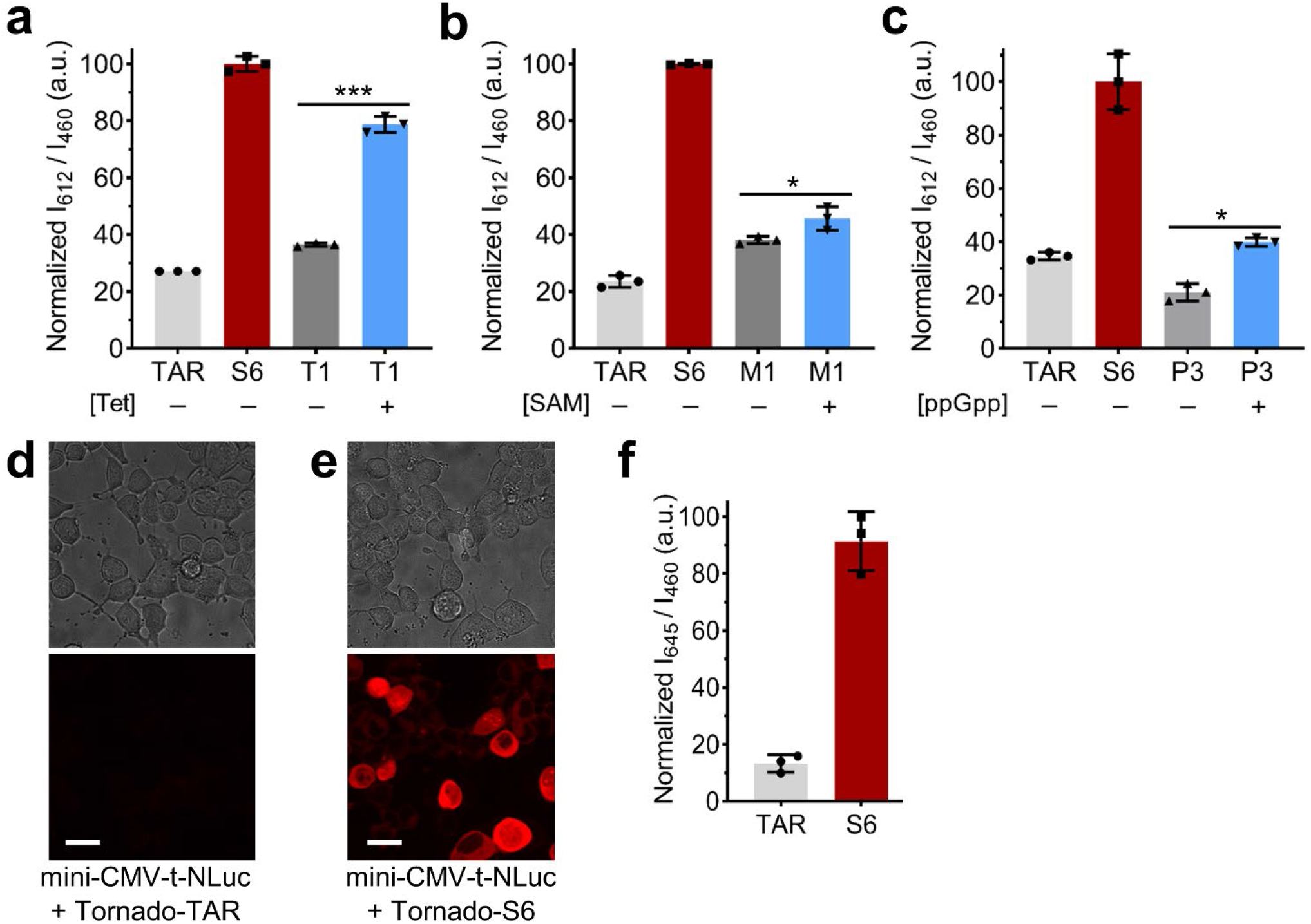
(**a**) I_612_/I_460_ ratiometric BRET signals of genetically encoded t-NLuc/T1 sensors in BL21 Star™ (DE3) cell lysates in the absence or presence of additional 100 μM tetracycline. (**b**) I_612_/I_460_ ratiometric BRET signals of genetically encoded t-NLuc/M1 sensors in BL21 Star™ (DE3) cell lysates in the absence or presence of additional 1 mM SAM. (**c**) I_612_/I_460_ ratiometric BRET signals of genetically encoded t-NLuc/P3 sensors in BL21 Star™ (DE3) cell lysates in the absence or presence of additional 100 μM ppGpp. t-NLuc/TAR (without the Pepper unit) acted as a negative control, and the t-NLuc/S6 construct acted as a positive control. The measurement was performed at 25°C in the presence of 5 μM HBC620 right after adding 1% (v/v) furimazine substrate. (**d, e**) Bright field and fluorescence channel imaging of HEK-293T cells at 24 h after transfection with (**d**) mini-CMV-t-NLuc and pAV-U6+27-Tornado-TAR or with (**e**) mini-CMV-t-NLuc and pAV-U6+27-Tornado-S6. Fluorescence images were collected through a 593/20 nm filter after irradiation with a 561 nm laser at 25°C in the presence of 5 μM HBC620. Scale bar, 20 μm. (**f**) The I_645_/ I_460_ ratiometric signals of HEK-293T cells after confirming the successful transfection with mini-CMV-t-NLuc and pAV-U6+27-Tornado-TAR or pAV-U6+27-Tornado-S6. The measurement was performed in a plate reader at 25°C in the presence of 5 μM HBC620 at 30 s after adding 1% (v/v) furimazine substrate. Shown are the mean and SD values from three independent replicates. *p< 0.05, ***p< 0.001 in two-tailed student’s t-test.

Lastly, we also tested if our RNA-based BRET system can also be genetically encoded and function within mammalian cells. For this purpose, we transfected our S6 construct into HEK-293T cells using a previously reported “Tornado” expression system.^66^ This Tornado platform enables rapid intracellular RNA circularization and protects RNA inserts from exonuclease degradation to increase the cellular expression levels of RNA sensors. After validating the expression of sensors in the transfected HEK-293T cells (Fig. 4d and 4e), the I_645_/I_460_ ratiometric signals were compared between t-NLuc/TAR- and t-NLuc/S6-expressing cells. An ∼6.9-fold increase in the BRET signal was observed in cells expressing t-NLuc/S6 (Fig. 4f), demonstrating the feasibility and potential function of this RNA-based BRET system in living mammalian cells.

## Conclusions

In this project, we developed the first genetically encoded RNA-based BRET sensors, which can be used for intracellular detection and imaging of different target analytes. On the basis of natural RNA–protein interactions, a luciferase enzyme donor and an FR aptamer acceptor was coupled together to form an effective BRET platform. Modular BRET sensors have been further engineered to detect antibiotics, metabolites and signaling molecules in both living cells and cell lysates. Following similar design principles, we expect the potential development of modular bioluminescent sensors for a wide range of target species, including small molecules, ions, RNAs, and proteins.

Another potential application of these BRET sensors is for the study of cellular RNA–protein interactions. By replacing TAR–Tat with other RNA–protein pair of interest, the BRET signals can function as a genetically encoded indicator to determine the strength and distribution of RNA–protein interactions in living biological system. Meanwhile, the potential usage of these RNA-based BRET sensors for *in vivo* imaging and high-throughput screening of RNA-targeted drugs can also be anticipated. Indeed, these general and modular BRET sensors can be a great complementary tool to the existing fluorescent RNA assays.

On the other hand, we have to mention that the current dynamic range and efficiency of these RNA-based BRET sensors are still quite limited. More systematic studies are still needed to further optimize FR/dye and luciferase/substrate pairs with better matched wavelength, brightness, and relative orientation. Nevertheless, this current study has now opened the door for new functions of RNA molecules and nanodevices. A wide variety of design and applications of RNA sensors can be expected in this emerging field.

## Data availability

The data found in this study are available within the main text and in the ESI online.

## Author contributions

Q. Y., K. R., and M. Y. designed the experiments. L. M. and Q. Y. performed most majority of the experiments and analyzed the data. A. P. K. K. K. synthesized and purified the t-NLuc protein. R. W., Z. S., and R. Z. synthesized and characterized some RNA sensors. K. R. and M. Y. supervised the whole project. L. M. and M. Y. wrote the manuscript. All the authors discussed the result shown in the manuscript.

## Acknowledgements

The authors gratefully acknowledge the support from an NSF CAREER award, Alfred P. Sloan Research Fellowship, Camille Dreyfus Teacher-Scholar Award and UMass Amherst start-up grant to M. You. Q. Yu and R. Zheng were also supported by NIH T32GM008515. We are grateful to Dr. James Chambers for the assistance in fluorescence imaging. The miniCMV-Nluc-tDeg plasmid was a gift from Prof. Samie R. Jaffrey. The authors also thank other members in the You Lab for useful discussion and valuable comments.

## Notes and references

1. E. C. Greenwald, S. Mehta and J. Zhang, Chem. Rev., 2018, 24, 11707–11794.

2. A. J. Syed and J. C. Anderson, Chem. Soc. Rev., 2021, 50, 5668–5705.

3. T. Wilson and J. W. Hastings, Curr. Rev. Cell Dev. Biol., 1998, 14, 197–230.

4. T. Ozawa, H. Yoshimura and S. B. Kim, Anal. Chem., 2013, 85, 590–609.

5. X. Zhou, S. Mehta and J. Zhang, Trends Biochem. Sci., 2020, 45, 889–905.

6. M. Tantama, Y. P. Hung and G. Yellen, J. Am. Chem. Soc., 2011, 133, 10034–10037.

7. A. B. Dippel, W. A. Anderson, J. H. Park, F. H. Yildiz and M. C. Hammond, ACS Chem. Biol., 2020, 15, 904–914.

8. T. W. Chen, T. J. Wardill, Y. Sun, S. R. Pulver, S. L. Renninger, A. Baohan, E. R. Schreiter, R. A. Kerr, M. B. Orger, V. Jayaraman, L. L. Looger, K. Svoboda and D. S. Kim, Nature, 2013, 499, 295–300.

9. R. Griss, A. Schena, L. Reymond, L. Patiny, D. Werner, C. E. Tinberg, D. Baker and K. Johnsson, Nat. Chem. Biol., 2014, 10, 598–603.

10. Z. Sun, T. Nguyen, K. McAuliffe and M. You, Nanomaterials (Basel), 2019, 9, 233.

11. Q. Yu, K. Ren and M. You, Nanoscale, 2021, 13, 7988–8003.

12. Y. Su and M. C. Hammond, Curr. Opin. Biotechnol., 2020, 63, 157–166.

13. J. S. Paige, T. Nguyen-Duc, W. Song and S. R. Jaffrey, Science, 2012, 335, 1194.

14. C. A. Kellenberger, C. Chen, A. T. Whiteley, D. A. Portnoy and M. C. Hammond, J. Am. Chem. Soc., 2015, 137, 6432–6435.

15. J. Wu, S. Zaccara, D. Khuperkar, H. Kim, M. E. Tanenbaum and S. R. Jaffrey, Nat. Methods, 2019, 16, 862–865.

16. R. Griss, A. Schena, L. Reymond, L. Patiny, D. Werner, C. E. Tinberg, D. Baker and K. Johnsson, Nat. Chem. Biol., 2014, 10, 598–603.

17. K. Tenda, B. van Gerven, R. Arts, Y. Hiruta, M. Merkx and D. Citterio, Angew. Chem. Int. Ed. Engl, 2018, 57, 15369–15373.

18. S. Liu, Y. Su, M. Z. Lin and J. A. Ronald, ACS Chem. Biol., 2021, 16, 2707–2718.

19. Y. Su, J. R. Walker, Y. Park, T. P. Smith, L. X. Liu, M. P. Hall, L. Labanieh, R. Hurst, D. C. Wang, L. P. Encell, N. Kim, F. Zhang, M. A. Kay, K. M. Casey, R. G. Majzner, J. R. Cochran, C. L. Mackall, T. A. Kirkland and M. Z. Lin, Nat. Methods, 2020, 17, 852–860.

20. M. P. Hall, J. Unch, B. F. Binkowski, M. P. Valley, B. L. Butler, M. G. Wood, P. Otto, K. Zimmerman, G. Vidugiris, T. Machleidt, M. B. Robers, H. A. Benink, C. T. Eggers, M. R. Slater, P. L. Meisenheimer, D. H. Klaubert, F. Fan, L. P. Encell and K. V. Wood, ACS Chem. Biol., 2012, 7, 1848–1857.

21. C. G. England, E. B. Ehlerding and W. Cai, Bioconjug. Chem., 2016, 27, 1175–1187.

22. F. Bouhedda, A. Autour and M. Ryckelynck, Int. J. Mol. Sci., 2017, 19, 44.

23. J. S. Paige, K. Y. Wu and S. R. Jaffrey, Science, 2011, 333, 642–646.

24. W. Song, R. L. Strack and S.R. Jaffrey, Nat. Methods., 2013, 10, 873–878.

25. C. A. Kellenberger, S. C. Wilson and M. C. Hammond, J. Am. Chem. Soc., 2013, 135, 4906–4909.

26. M. You, J. L. Litke and S. R. Jaffrey, Proc. Natl. Acad. Sci. USA., 2015, 112, E5765–E5765.

27. E. B. Porter, J. T. Polaski, M. M. Morck and R. T. Batey, Nat. Chem. Biol., 2017 13, 295–301.

28. X. Li, J. L. Litke, S. K. Dey, S. R. Suter and S. R. Jaffrey, J. Am. Chem. Soc., 2020, 142, 14117–14124.

29. H. Kim and S. R. Jaffrey, Cell Chem. Biol., 2019, 26, 1725–1731.

30. Y. Xu, D. W. Piston and C. H. Johnson, Proc. Natl. Acad. Sci. USA., 1999, 96, 151–156.

31. K. D. G. Pfleger, K. A. Eidne, Nat. Methods., 2006, 3, 165–175.

32. X. Xu, M. Soutto, Q. Xie, S. Servick, C. Subramanian, A. G. von Arnim and C. H. Johnson, Proc. Natl. Acad. Sci. USA., 2007, 104, 10264–10269.

33. A. Dragulescu-Andrasi, C. T. Chan, T. F. Massoud and S. S. Gambhir, Proc. Natl. Acad. Sci. USA., 2011, 108, 12060–12065.

34. J.D. Puglisi, L. Chen, S. Blanchard and A. D. Frankel, Science, 1995, 270, 1200–1203.

35. X. Ye, R. A. Kumar and D. J. Patel, Chem. Biol., 1995, 2, 827–840.

36. N. L. Greenbaum, Structure 1996, 4, 5–9.

37. J. Karn, J. Mol. Biol., 1999, 293, 235–254.

38. M. J. Selby, E. S. Bain, P. A. Luciw and B. M. Peterlin, Genes & Dev, 1989, 3, 547–558.

39. I. D’Orso and A. D. Frankel, Nat. Struct. Mol. Biol., 2010, 17, 815–821.

40. J. D. Puglisi, S. Blanchard and A. D. Frankel, Science, 1995, 270, 1200–1203.

41. S. Roy, C. Chen, C. A. Rosen, and N. Sonenberg, Genes & Dev, 1990, 4, 1365–1373.

42. G. S. Filonov and S. R. Jaffrey, J. Am. Chem. Soc., 2014, 136, 16299–16308.

43. R. J. Trachman, N. A. Demeshkina, M. W. L. Lau, S. S. S. Panchapakesan, S. C. Y. Jeng, P. J. Unrau and A. R. Ferre-D’Amare, Nat. Chem. Biol., 2017, 13, 807–813.

44. E. V. Dolgosheina, S. C. Jeng, S. S. Panchapakesan, R. Cojocaru, P. S. Chen, P. D. Wilson, N. Hawkins, P. A. Wiggins and P. J. Unrau, ACS Chem. Biol., 2014, 9, 2412–2420.

45. X. Chen, D. Zhang, N. Su, B. Bao, X. Xie, F. Zuo, L. Yang, H. Wang, L. Jiang, Q. Lin, M. Fang, N. Li, X. Hua, Z. Chen, C. Bao, J. Xu, W. Du, L. Zhang, Y. Zhao, L. Zhu, J. Loscalzo and Y. Yang, Nat. Biotechnol., 2019, 37, 1287–1293.

46. K. Huang, X. Chen, C. Li, Q. Song, H. Li, L. Zhu, Y. Yang and A. Ren, Nat. Chem. Biol., 2021, 17, 1289–1295.

47. U. Schulze-Gahmen and J. H. Hurley, Proc. Natl. Acad. Sci. USA., 2018, 115, 12973–12978.

48. A. M. P. Romani, Arch. Biochem. Biophys., 2011, 512, 1–23.

49. W. Jahnen-Dechent and M. Ketteler, Clin. Kidney J., 2012, 5, 3–14.

50. B. S. Speer, N. B. Shoemaker and A. A. Salyers, Clin. Microbiol. Rev., 1992, 5, 387–399.

51. A. Wittmann and B. Suess, Mol. Biosyst., 2011, 7, 2419–2427.

52. M. Zuker, Nucleic Acids Res., 2003, 31, 3406–3415.

53. N. E. Holmes and P. G. P. Charles, Clin. Med. Ther., 2009, 1, 471–482.

54. S. C. Lu, Int. J. Biochem. Cell Biol., 2000, 32, 391–395.

55. G. I. Papakostas, J. E. Alpert and M. Fava, Curr. Psychiatry Rep., 2003, 5, 460–466.

56. J. Abranches, A. R. Martinez, J. K. Kajfasz, V. Chavez, D. A. Garsin and J. A. Lemos, J. Bacteriol., 2009, 7, 2248–2256.

57. W. A. Haseltine and R. Block, Proc. Natl. Acad. Sci. USA., 1973, 70, 1564–1568.

58. J. Gallant, L. Palmer and C. C. Pao, Cell, 1977, 11, 181–185.

59. E. Loh, O. Dussurget, J. Gripenland, K. Vaitkevicius, T. Tiensuu, P. Mandin, F. Repoila, C. Buchrieser, P. Cossart and J. Johansson, Cell, 2009, 139, 770–779.

60. M. E. Sherlock, N. Sudarsan and R. R. Breaker, Proc. Natl. Acad. Sci. USA., 2018, 115, 6052–6057.

61. A. Peselis and A. Serganov, Nat. Chem. Biol., 2018, 14, 887–894.

62. Z. Sun, R. Wu, B. Zhao, R. Zeinert, P. Chien and M. You, Angew. Chem. Int. Ed. Engl., 2021, 60, 24070–24074.

63. J. Chu, J. Qian, Y. Zhuang, S. Zhang and Y. Li, Appl. Microbiol. Biotechnol., 2013, 97, 41–49.

64. V. J. Hernandez and H. Bremer, J. Biol. Chem., 1990, 265, 11605–11614.

65. L. I. Pizer and J. P. Merlie, J. Bacteriol., 1973, 114, 980–987.

66. J. L. Litke and S. R. Jaffrey, Nat. Biotechnol., 2019, 37, 667–675.

