## Supplementary Information for "Genetically Encoded RNA-based Bioluminescence Resonance Energy Transfer (BRET) Sensors"

#### Contents

##### Supplementary Figures

**Fig. S1**. Fluorescence signals and bioluminescence spectra of the t-NLuc/TAR-Pepper construct.

**Fig. S2**. Effect of incubation time and substrate concentration on the t-NLuc/S6 BRET signal.

**Fig. S3**. Effect of $Mg^{2+}$ concentration on the t-NLuc/S6 BRET signal.

**Fig. S4**. Effect of buffer condition on the BRET and fluorescence signal of the t-NLuc/S6 construct.

**Fig. S5**. Fluorescence and BRET signals of different tetracycline, SAM, and ppGpp sensors

**Fig. S6**. Effect of induction time and promotor on the cellular BRET signal.

##### Supplementary Table

**Table S1**. RNA sequences used in this work.

### 1. Supplementary Figures

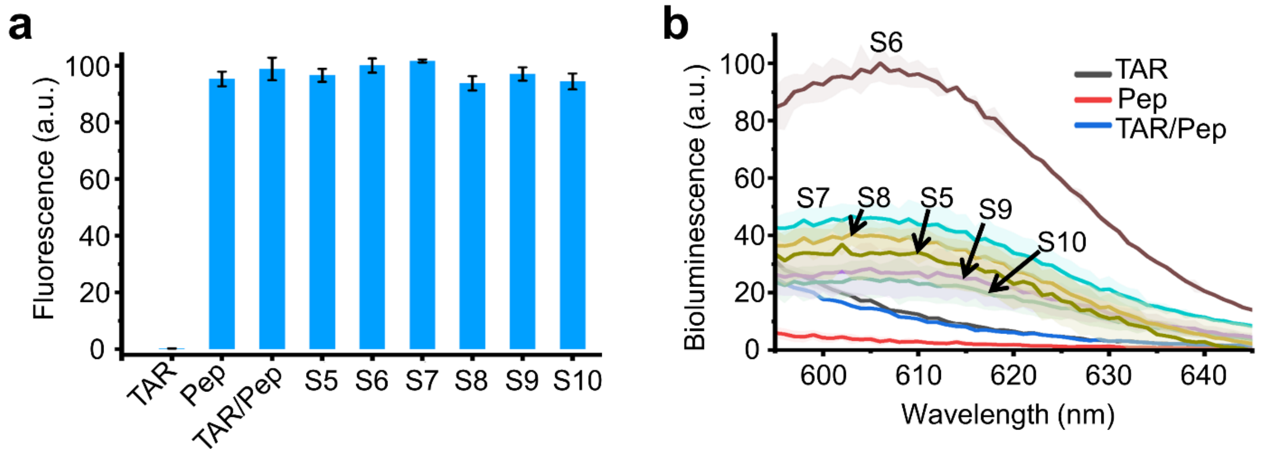


**Fig. S1.** (**a**) Fluorescence intensities were measured at 612 nm after excitation at 572 nm using a solution containing 100 nM t-NLuc, 1 μM RNA, and 5 μM HBC620 at 25°C. S5–S10 represent the t-NLuc/TAR-Pepper construct containing 5–10-base pair-long linkers. Without the Pepper unit, the mixture of t-NLuc with TAR was used as a negative control. Without conjugating with TAR, Pepper acted as a positive control. (**b**) Bioluminescence spectra of the t-NLuc/TAR-Pepper construct containing 5–10-base-pair-long linkers (S5–S10). The measurement was performed in a solution containing 100 nM t-NLuc, 1 μM RNA, and 5 μM HBC620 at 25°C. The mixture of t-NLuc with TAR (without Pepper) or Pepper (without TAR conjugation) were used as negative controls. Shown are the mean and standard deviation (SD) values (error bar or shaded area) from ≥three independent replicates.

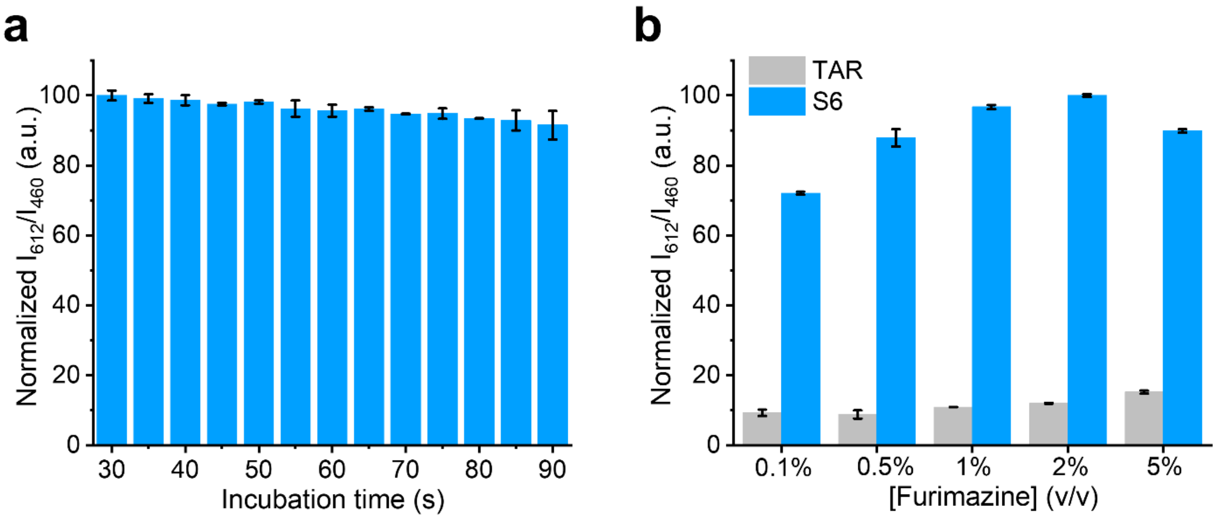


**Fig. S2.** (**a**) The $I_{612}/I_{460}$ (acceptor/donor) ratiometric signal of t-NLuc/S6 as measured at different time point after adding 1% (v/v) furimazine substrate at 30 s. (**b**) The $I_{612}/I_{460}$ ratiometric signal as measured at 1 min after adding different amount of the furimazine substrate. The measurements were performed in a solution containing 100 nM t-NLuc, 1 μM S6 or TAR, and 5 μM HBC620 at 25°C. Shown are the mean and the standard error of the mean (SEM) values from three independent replicates.

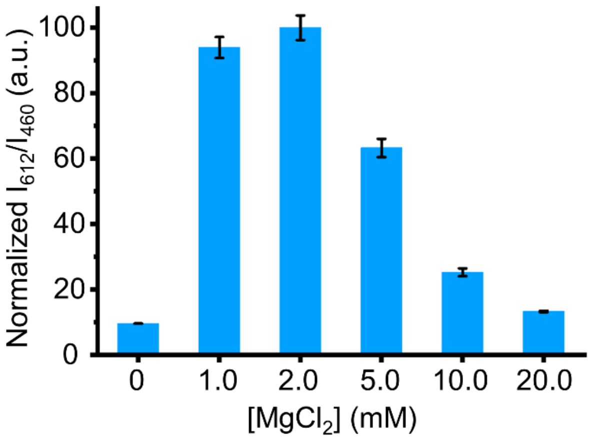


**Fig. S3.** The effect of $Mg^{2+}$ concentration on the $I_{612}/I_{460}$ signal ratio of the t-NLuc/S6 construct. The measurements were performed in a solution containing 100 nM t-NLuc, 1 μM S6, and 5 μM HBC620 at 25°C. Shown are the mean and SEM values from three independent replicates.

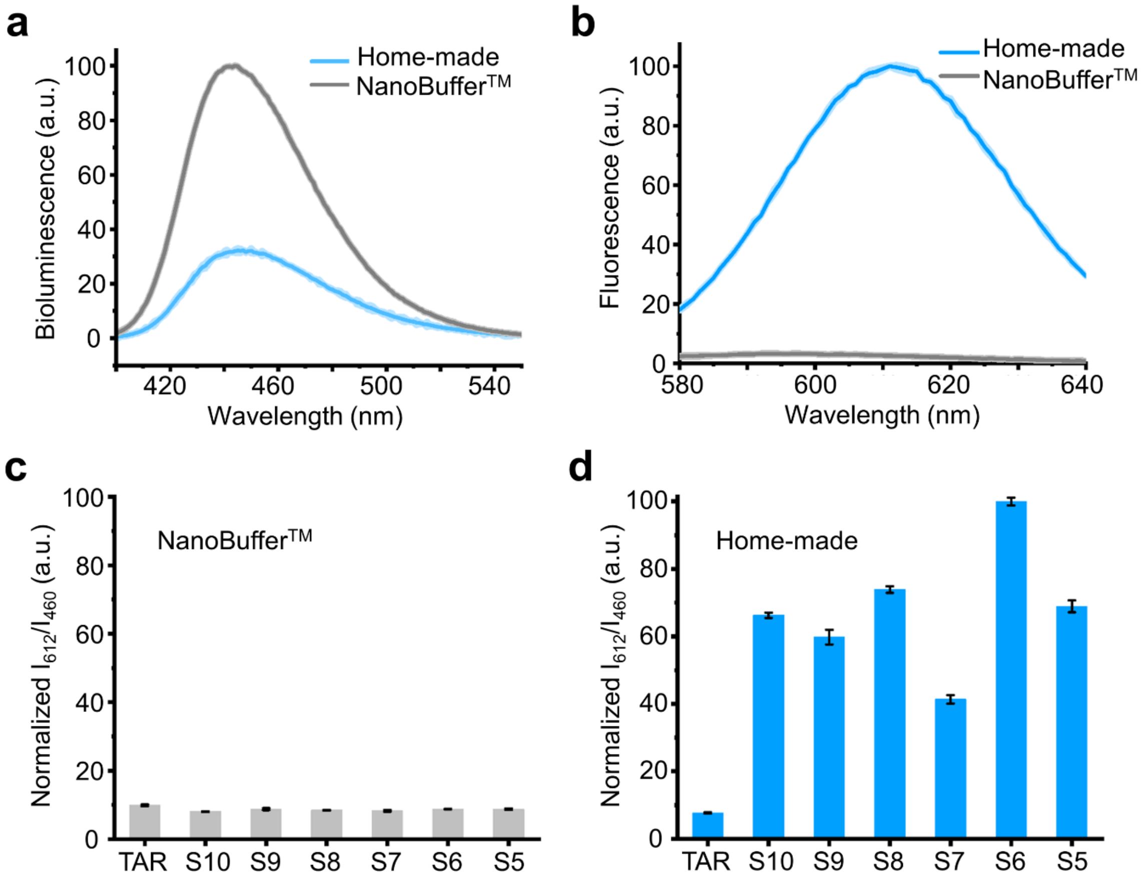


**Fig. S4.** The effect of buffer condition on the (**a**) bioluminescence spectrum of t-NLuc/S6 and (**b**) fluorescence spectrum of Pepper after excitation at 572 nm. The home-made buffer consists of 40 mM HEPES, 100 mM KCl, and 0.1% DMSO at pH=7.5. (**c**, **d**) The $I_{612}/I_{460}$ ratiometric signal of the t-NLuc/TAR-Pepper construct containing 5–10-base pair-long linkers was measured in either NanoBuffer™ or home-made buffer, both of which contain 2 mM $MgCl_2$. The mixture of t-NLuc with TAR (without Pepper) was used as a negative control. The measurements were performed in a solution containing 100 nM t-NLuc, 1 μM RNA, and 5 μM HBC620 at 25°C. Shown are the mean and SEM values (error bar or shaded area) from three independent replicates.

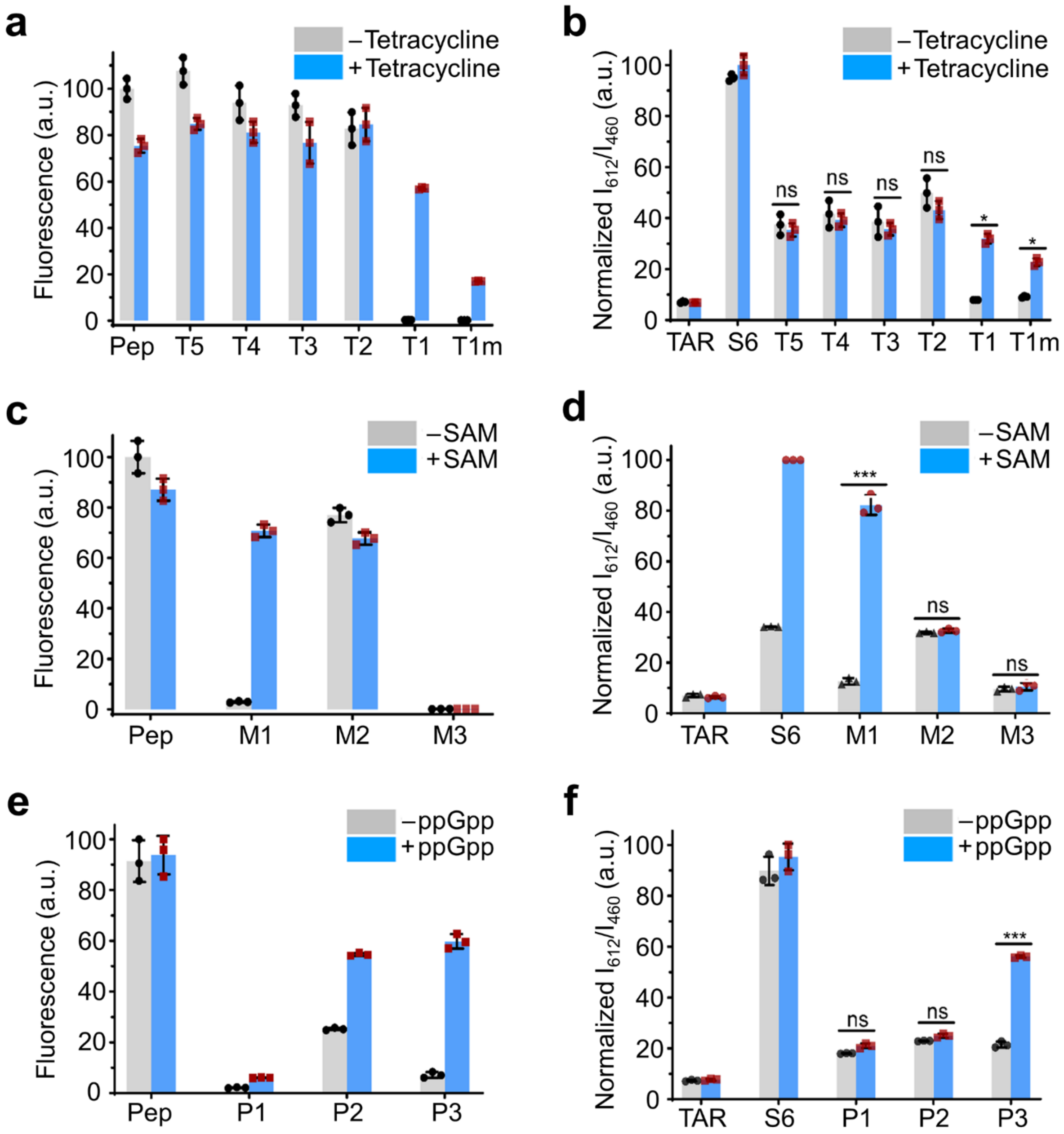


**Fig. S5.** (**a**, **c**, **e**) Fluorescence intensities of different Pepper-based tetracycline, SAM, and ppGpp sensors as measured at 612 nm upon excitation at 572 nm. The measurements were performed in a solution containing 100 nM t-NLuc, 1 μM RNA, 5 μM HBC620, and in the absence or presence of 100 μM corresponding target at 25°C. The T1m construct contains one base pair mutation different from the T1 RNA. The t-NLuc/T1, t-NLuc/M1 and t-NLuc/P3 sensors were identified as the optimal system that exhibited the highest fluorescence enhancement. (**b**, **d**, **f**) The $I_{612}/I_{460}$ ratiometric BRET signal of different Pepper-based tetracycline, SAM, and ppGpp sensors as measured in the absence or presence of 100 μM corresponding target. The mixture of t-NLuc with TAR or S6 acted as a negative control or a positive control, respectively. The measurements were performed in a solution containing 100 nM t-NLuc, 1 μM RNA, 5 μM HBC620 at 25°C after adding 1% (v/v) furimazine substrate. Shown are the mean and SD values from three independent replicates in each case. *$p < 0.05$, ***$p < 0.001$ in two-tailed student's t-test; ns, not significant.

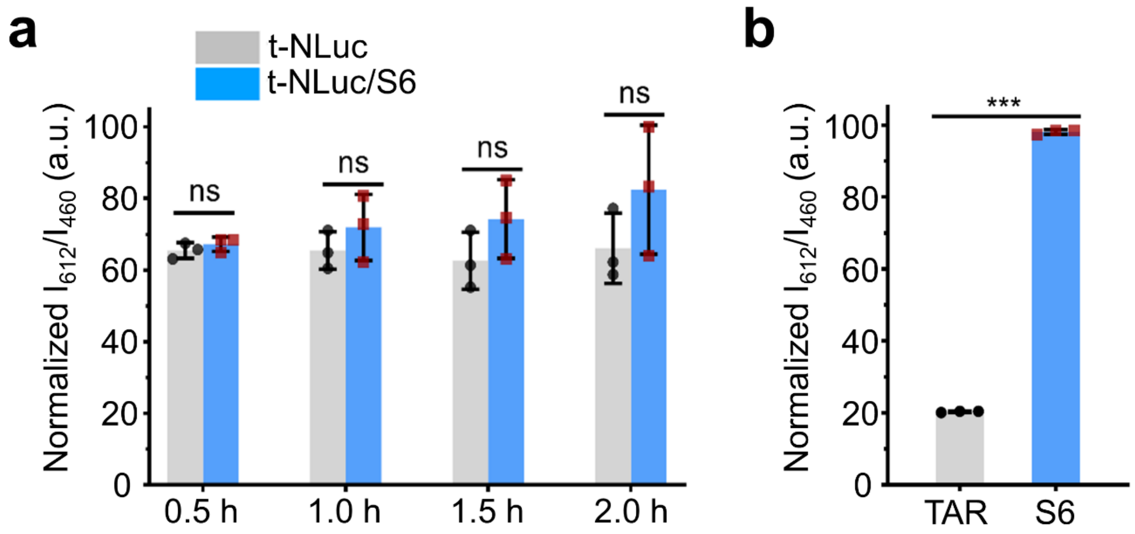


**Fig. S6.** (**a**) $I_{612}/I_{460}$ ratiometric BRET signal as obtained in BL21 Star™ (DE3) cells that express only the t-NLuc or the t-NLuc/S6 construct after different time of IPTG induction. The expression of S6 RNA and t-NLuc protein were both under T7 promoters. (**b**) $I_{612}/I_{460}$ ratiometric BRET signal as measured in BL21 Star™ (DE3) cells that express t-NLuc under the control of only lac promotor. The RNA construct was always expressed under a T7 promoter after 2 h of IPTG induction. Shown are the mean and SD values from three independent replicates. ***p< 0.001 in two-tailed student's t-test; ns, not significant.

#### 2. Supplementary Table

**Table S1. RNA sequences used in this work.** Blue: target binding aptamer; gray: transducer; orange: TAR RNA; red: Pepper RNA; green: Broccoli RNA; yellow: Mango II RNA.

| Name | RNA sequence (5' to 3') |
|---|---|
| TAR | GGCUCGUUGAGCUCAUUAGCUCCGAGCC |
| Pepper | GAUGAUCCCCAAUCGUGGCGUGUCGGCCUGCUUCGGCAGGCACUGGCGCCGGGAUCAUC |
| S5 | GGAGGCUCGUUGAGCCCAAUCGUGGCGUGUCGGCCUGCUUCGGCAGGCACUGGCGCCGGCUCCGAGCCUCC |
| S6 | GGAGGCUCGUUGAGCUCCAAUCGUGGCGUGUCGGCCUGCUUCGGCAGGCACUGGCGCCGAGCUCCGAGCCUCC |
| S7 | GGAGGCUCGUUGAGCUUCCAAUCGUGGCGUGUCGGCCUGCUUCGGCAGGCACUGGCGCCGAAGCUCCGAGCCUCC |
| S8 | GGAGGCUCGUUGAGCUGUCCAAUCGUGGCGUGUCGGCCUGCUUCGGCAGGCACUGGCGCCGACAGCUCCGAGCCUCC |
| S9 | GGAGGCUCGUUGAGCUGGUCCAAUCGUGGCGUGUCGGCCUGCUUCGGCAGGCACUGGCGCCGACCAGCUCCGAGCCUCC |
| S10 | GGAGGCUCGUUGAGCUGGGUCCAAUCGUGGCGUGUCGGCCUGCUUCGGCAGGCACUGGCGCCGACCCAGCUCCGAGCCUCC |
| Broccoli | GAGAGGAGACGGUCGGGUCCAGAUAUUCGUAUCUGUCGAGUAGAGUGUGGGCUCCUCUC |
| B1 | GAGAGGAGACGGUCGGGUCCAGAUAGGCUCGUUGAGCUCAUUAGCUCCGAGCCUAUCUGUCGAGUAGAGUGUGGGCUCCUCUC |
| B2 | GAGAGGAGACGGUCGGGUCCAGACUCGUUGAGCUCAUUAGCUCCGAGUCUGUCGAGUAGAGUGUGGGCUCCUCUC |
| B3 | GAGAGGAGACGGUCGGGUCCACUCGUUGAGCUCAUUAGCUCCGAGUGUCGAGUAGAGUGUGGGCUCCUCUC |
| Mango II | GCGUACGAAGGAGAGGAGAGGAAGAGGAGAGUACGC |
| Ma1 | GGAGGCUCGUUGAGCUGCGUACGAAGGAGAGGAGAGGAAGAGGAGAGUACGCAGCUCCGAGCCUCC |
| Ma2 | GGAGGCUCGUUGAGCUGUACGAAGGAGAGGAGAGGAAGAGGAGAGUACAGCUCCGAGCCUCC |
| Ma3 | GGAGGCUCGUUGAGCUGCGCGUACGAAGGAGAGGAGAGGAAGAGGAGAGUACGCGCAGCUCCGAGCCUCC |
| T1 | GGAGGCUCGUUGAGCUCCAAUCGUGGCGUGUCGGAAAACAUACCAGAUUUCGAUCUGGAGAGGUGAAGAAUACGACCACCUCACUGGCGCCGAGCUCCGAGCCUCC |
| T1m | GGAGGCUCGUUGAGCUCCAAUCGUGGCGUGUCGAAAAACAUACCAGAUUUCGAUCUGGAGAGGUGAAGAAUACGACCACCUUACUGGCGCCGAGCUCCGAGCCUCC |
| T2 | GGAGGCUCGUUGAGCUCCAAUCGUGGCGUGUCGGCAAAACAUACCAGAUUUCGAUCUGGAGAGGUGAAGAAUACGACCACCUGCACUGGCGCCGAGCUCCGAGCCUCC |
| T3 | GGAGGCUCGUUGAGCUCCAAUCGUGGCGUGUCGGCCAAAACAUACCAGAUUUCGAUCUGGAGAGGUGAAGAAUACGACCACCUGGCACUGGCGCCGAGCUCCGAGCCUCC |
| T4 | GGAGGCUCGUUGAGCUCCAAUCGUGGCGUGUCGGCCUAAAACAUACCAGAUUUCGAUCUGGAGAGGUGAAGAAUACGACCACCUAGGCACUGGCGCCGAGCUCCGAGCCUCC |

| | |
|---|---|
| T5 | GGAGGCUCGUUGAGCUCCAAUCGUGGCGUGUCGGCCUGAAAACAUACCAGAUUUCGAUCUGGAGAGGUGAAGAAUACGACCACCUCAGGCACUGGCGCCGAGCUCCGAGCCUCC |
| M1 | GGAGGCUCGUUGAGCUCCAAUCGUGGCGUGUCGGCGAAAGGAUGGCGGAAACGCCAGAUGCCUUGUAACCGAAAGGGCACUGGCGCCGAGCUCCGAGCCUCC |
| M2 | GGAGGCUCGUUGAGCUCCAAUCGUGGCGUGUCGUUCCCGAAAGGAUGGCGGAAACGCCAGAUGCCUUGUAACCGAAAGGGGGAAACUGGCGCCGAGCUCCGAGCCUCC |
| M3 | GGAGGCUCGUUGAGCUCCAAUCGUGGCGUGUCGAAGAAAGGAUGGCGGAAACGCCAGAUGCCUUGUAACCGAAAGGUUACUGGCGCCGAGCUCCGAGCCUCC |
| P1 | CAGCGACCGAGCGGUACAAUGACUGGCGCCGAGCUCCGAGCCUCCGCAAGGAGGCUCGUUGAGCUCCAAUCGUGGCGUGUCGCAACACCGUGAGCAUAAAAGGCUCA |
| P2 | CAGCGACCGAGCGGUACAAUCGACUGGCGCCGAGCUCCGAGCCUCCGCAAGGAGGCUCGUUGAGCUCCAAUCGUGGCGUGUCGCGAACACCGUGAGCAUAAAAGGCUCCA |
| P3 | CAGCGACCGAGCGGUACAAUCCCACUGGCGCCGAGCUCCGAGCCUCCGCAAGGAGGCUCGUUGAGCUCCAAUCGUGGCGUGUCGGGGAACACCGUGAGCAUAAAAGGCUCCA |

#### Reference


1. J. Wu, S. Zaccara, D. Khuperkar, H. Kim, M. E. Tanenbaum and S. R. Jaffrey, *Nat. Methods*, 2019, **16**, 862-865.
